# Therapeutic anti-IgE antibodies generate distinct functional immune complex signatures

**DOI:** 10.64898/2026.11.06.764804

**Authors:** María Miró Navarro, Alexander G. Geiger, Simona Notova, Andreas Frutiger, Martin F. Bachmann, Paul Engeroff

**Author notes:** equal contribution (María Miró Navarro, Alexander G. Geiger).

## Abstract

Therapeutic IgG anti-IgE antibodies are benchmarked by their ability to interfere with IgE–FcεRI binding, but their activity within IgE and IgG-containing immune complexes remains poorly defined. Here we show that omalizumab and ligelizumab form binary IgE–IgG and ternary allergen-containing immune complexes with distinct functional properties. In primary human leukocytes, these complexes exhibited antibody- and composition-dependent patterns of cellular association and activation. Using multiplexed focal molography, we resolved Fc receptor engagement across free and complexed antibody states and identified distinct receptor-binding signatures for omalizumab- and ligelizumab-containing complexes. Notably, allergen incorporation reversed the relative ability of the two antibodies to neutralize IgE–FcεRI interactions and remodelled FcγR binding. Thus, immune-complex composition is a determinant of therapeutic antibody function.

## Introduction

Therapeutic monoclonal antibodies have become a cornerstone of modern medicine through their ability to selectively target disease-associated molecules with high affinity^1^. A prominent example is the therapeutic targeting of immunoglobulin E (IgE) in allergic diseases, where the monoclonal antibody omalizumab has shown remarkable clinical efficacy^2–4^. Mechanistically, omalizumab interferes with the interaction between IgE and its high-affinity receptor FcεRI, which drives hypersensitivity responses in allergic effector cells such as basophils and mast cells^5–8^.

Ligelizumab was developed as a next-generation anti-IgE antibody with higher affinity for IgE and more potent IgE:FcεRI neutralization ability than omalizumab. However, despite these properties, ligelizumab did not demonstrate clinical superiority over omalizumab^9,10^. A possible explanation is the fact that ligelizumab is less potent at inhibiting interactions between IgE and the low-affinity IgE receptor CD23 (FcεRII) than omalizumab^10,11^. However, the contribution of CD23 to allergic pathology remains debated and has been challenged by some reports^12–14^. Thus, additional mechanisms may contribute to the functional differences between therapeutic anti-IgE antibodies.

IgG anti-IgE antibodies form “binary” IgE–IgG immune complexes (ICs), and allergen exposure may generate “ternary” IgE–IgG–allergen ICs, which may reshape engagement of both Fcε and Fcγ receptors^15–17^. FcγRI is the high-affinity IgG receptor, whereas FcγRIIa/b and FcγRIIIa/b are lower-affinity receptors whose engagement is favoured by multivalent ICs^18,19^. How IgE–IgG IC composition affects Fc receptor engagement and cellular activity remains unclear. Here, we compared omalizumab- and ligelizumab-containing ICs using a defined system based on the recombinant major cat allergen Fel d 1 and recombinant anti-Fel d 1 IgE in assays with primary human leukocytes and multiplexed focal molography^20,21^. We identify antibody- and composition-dependent patterns of Fc receptor engagement and cellular activation.

Strikingly, allergen incorporation reverses the relative ability of omalizumab and ligelizumab to neutralize IgE–FcεRI interactions and also shifted relative FcγR engagement, revealing IC signatures as an additional functional layer of therapeutic antibody activity.

## Results

### Ligelizumab and omalizumab ICs differentially modulate allergen association across leukocyte populations

Because Fcε and Fcγ receptors are differentially distributed across leukocyte populations, we hypothesized that IgE–IgG complex formation could alter the cellular distribution of IgE-associated allergen.

To visualize Fc receptor expression patterns across major human blood cell populations, we analysed a public scRNA-seq dataset (see methods) and performed flow cytometry analysis of whole blood cells from healthy human donors (Fig. 1a–c, Fig S1). As expected, FcεRI expression or surface-bound IgE were highest in basophils, but also detectable in monocytes, whereas FcεRII (CD23) expression was largely confined to B cells. FcγRI was expressed in monocytes, whereas low-affinity FcγRs were more broadly distributed across leukocytes. Fc receptors were largely absent from T cells.

**Figure 1.**
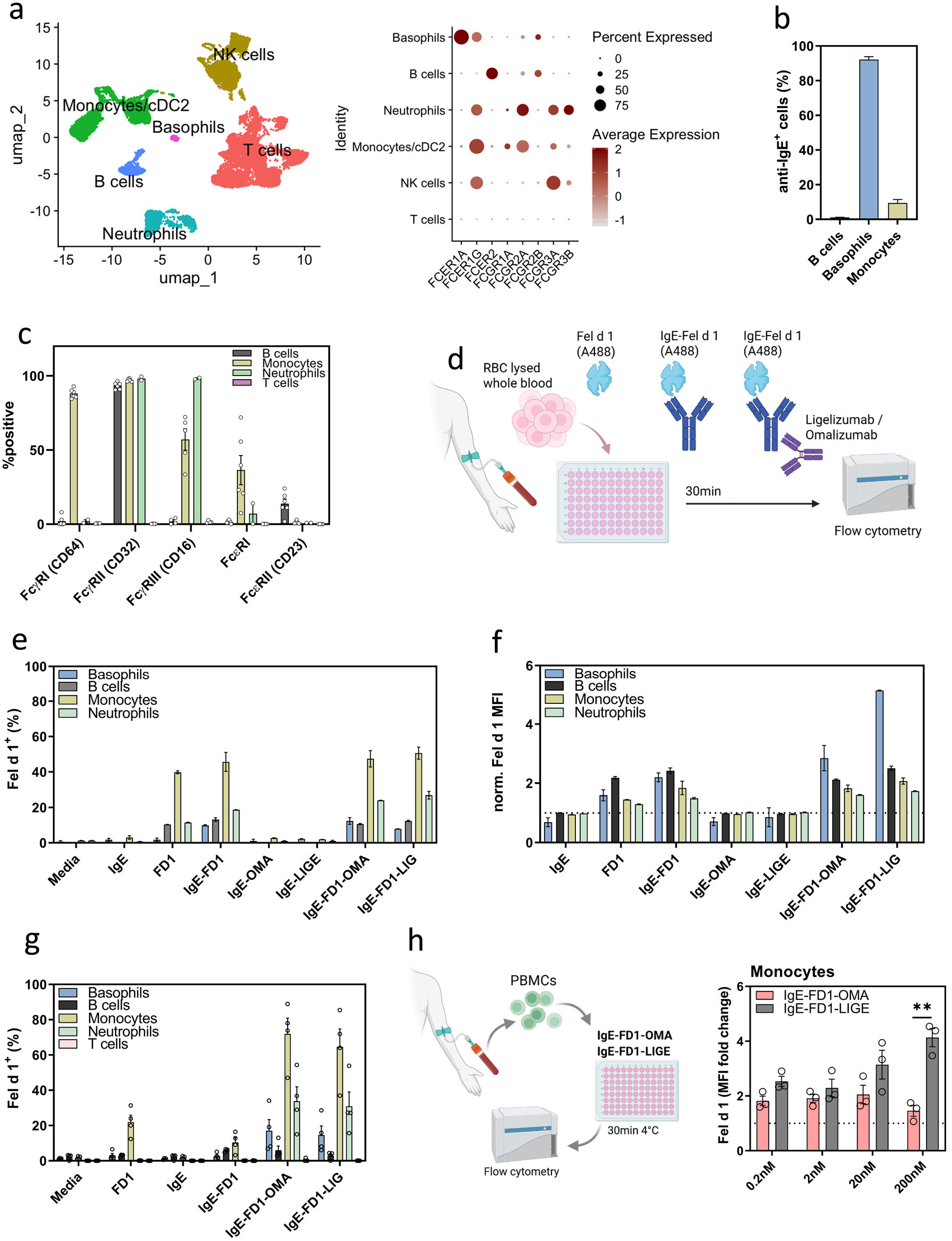
| Ligelizumab and omalizumab ICs differentially modulate allergen association across leukocyte populations. **a,** UMAP visualization of major leukocyte populations and expression of Fc receptor transcripts. **b,** surface-bound IgE on primary human leukocytes. **c,** Flow cytometric analysis of Fc receptor surface expression in healthy donors (n=2-6). **d–f,** Whole-blood leukocytes were incubated with fluorescent Fel d 1 alone or pre-formed ICs consisting of Fel d 1, human IgE, and omalizumab or ligelizumab before quantification of cell-associated allergen by flow cytometry. **d,** Experimental design. **e,** % Fel d 1□ cells at 4°C (one donor, technical replicates). **f,** Fel d 1 gMFI at 4°C (one donor, technical replicates). **g,** % Fel d 1□ cells at 37°C (n=4). **h,** Experimental design for PBMC titration experiments (n=3); Fel d 1 gMFI fold change normalized to medium control of CD14□ monocytes is shown. \*\**p* < 0.01 (Šídák’s multiple comparisons test). Symbols represent biological replicates; technical replicates were averaged per donor. Abbreviations used in figures: FD1 / FelD, Fel d 1; OMA, omalizumab; LIG / LIGE, ligelizumab; IC, immune complex.

To examine how therapeutic anti-IgE antibodies influence allergen association across these major cell types, whole-blood leukocytes were incubated with pre-formed ICs comprising fluorescently labelled Fel d 1, human IgE, and omalizumab or ligelizumab for 30 min at 4°C before quantifying cell-associated fluorescence by flow cytometry (Fig. 1d, Fig. S1). The allergen-containing ICs associated with multiple leukocyte populations, with monocytes showing the highest frequencies of Fel d 1□ cells (Fig. 1e). In basophils, we noted an increase in Fel d 1 MFI values in the presence of therapeutic anti-IgE antibodies, particularly for ligelizumab-containing ICs (Fig. 1f). At 37°C, both anti-IgE antibodies increased the percentage of Fel d 1 basophils, monocytes, and neutrophils, but not B cells, compared with Fel d 1 or IgE–Fel d 1 complexes alone (Fig. 1g). We next performed an experiment in peripheral blood mononuclear cells (PBMCs) at 4°C, where monocytes are enriched. In this setting, ligelizumab ICs produced a concentration-dependent increase (p = 0.0011) in monocyte association compared with omalizumab ICs (Fig. 1h, Fig. S2). This preferential monocyte association indicates that redistribution is governed not only by the introduction of an IgG Fc but also by the identity of the anti-IgE antibody.

Together, these findings show that anti-IgE introduction redistributes allergen-containing complexes across leukocyte populations in an antibody-dependent manner.

### Anti-IgE immune complexes induce cell-type- and antibody-dependent functional responses

To determine how cellular association translates into functional activation, we next investigated the capacity of ligelizumab- and omalizumab-containing ICs to activate immune cells.

Activation in whole blood was assessed by surface CD63 upregulation, a classical marker of FcεRI-mediated activation in basophils (Fig. 2a). Basophil responses to Fel d 1 or IgE– Fel d 1 alone were detectable but strongly donor-dependent (Fig. 2b): donors 1, 3, and 5 showed modest responses, donor 2 barely responded, but a strong response was observed in donor 4, who had elevated Fel d 1-specific IgE (0.462 kU_A_/L; Fig. S3). Pre-formed ICs activated basophils despite the presence of therapeutic anti-IgE antibody, and ternary ligelizumab-containing ICs gave the highest responses in each of the responding donors (Fig. 2b). In the most responsive donor, binary ICs showed a bell-shaped concentration dependence, declining at the highest antibody concentration. To compare conditions across donors despite these differences in curve shape and magnitude, we calculated the area under the concentration–response curve for each donor and condition (Fig. 2c). This revealed a composition-dependent reversal: in binary ICs, omalizumab-containing ICs gave higher basophil responses than ligelizumab-containing ICs, whereas in ternary ICs the relationship inverted, with ligelizumab-containing ICs the more activating (Fig. 2d). The reversal was present in all five donors (antibody × IC interaction, F(1,4) = 9.77, p = 0.035, two-way repeated-measures ANOVA), although neither IC class differed significantly from zero on its own.

**Figure 2.**
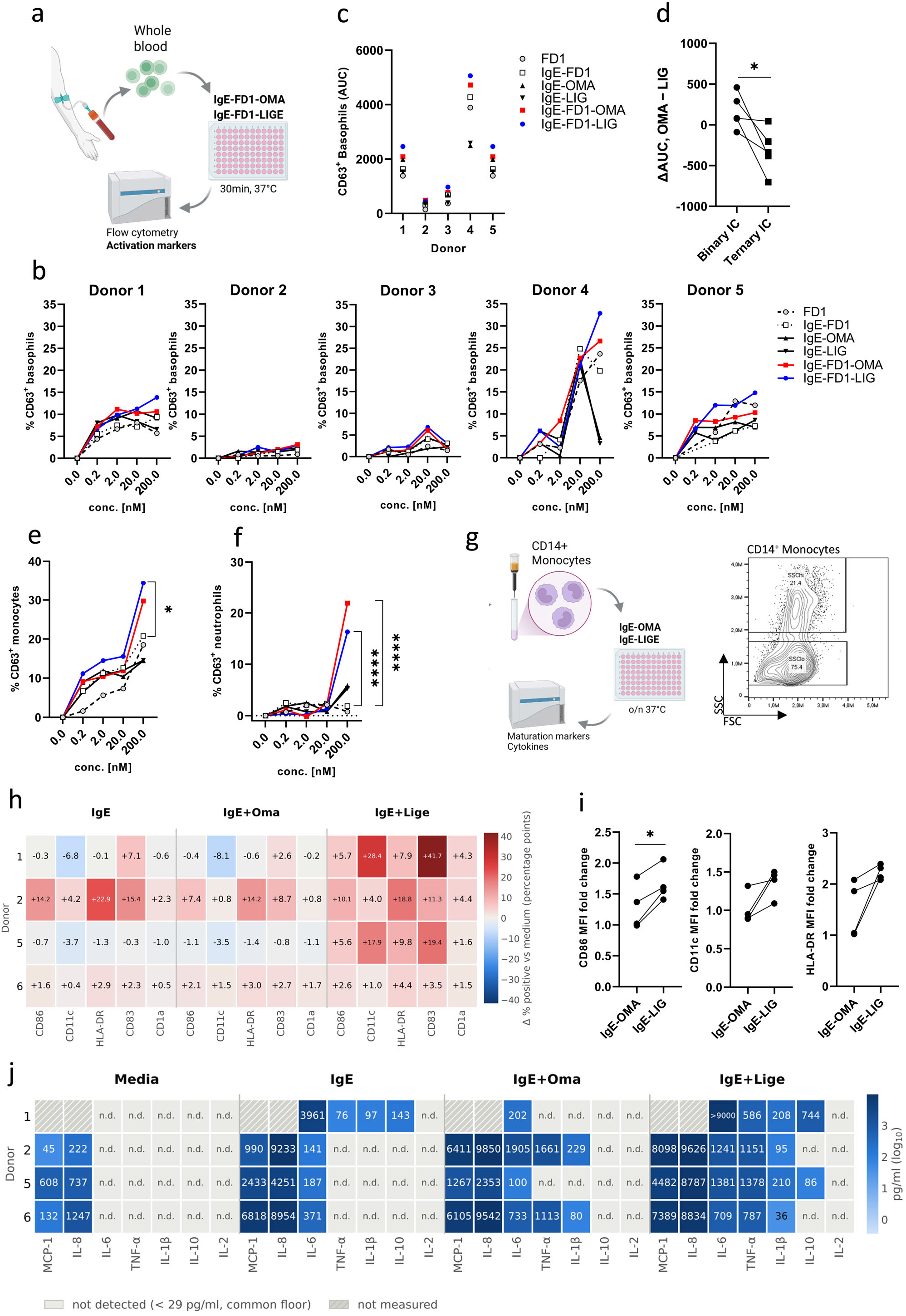
| Anti-IgE immune complexes induce cell-type- and antibody-dependent functional responses. **a**, Workflow for whole-blood stimulation with pre-formed binary (IgE–anti-IgE) and ternary (IgE–anti-IgE–Fel d 1) ICs, 30 min at 37 °C. **b**, CD63 upregulation on basophils across the Fel d 1 concentration range, shown for five individual donors. **c**, Area under the concentration–response curve (AUC) per donor and condition (n = 5). **d**, Within-donor difference in basophil AUC (omalizumab − ligelizumab) for binary and ternary ICs; lines connect paired values from the same donor (n = 5). *p = 0.035 for the antibody × IC interaction (two-way RM ANOVA). **e**, **f**, CD63 upregulation on monocytes (**e**) and neutrophils (**f**) in the same samples (mean values, n = 4). *p < 0.05, ****p < 0.0001 versus IgE–Fel d 1 (two-way RM ANOVA with Dunnett’s multiple comparisons test). **g**, Isolation and overnight stimulation of purified CD14 monocytes, with SSC gating. **h**–**j**, Monocytes stimulated with binary ICs (n = 4 donors). **h**, Change in the proportion of marker-positive SSC^hi^ cells relative to each donor’s own medium (percentage points). **i**, MFI fold change versus medium for CD86, CD11c, HLA-DR and CD83 in SSC^hi^ cells; lines connect paired values from the same donor. *p = 0.011 (two-tailed paired t-test). **j**, Cytokine concentrations in the same supernatants, shown as log□□ pg/ml; descriptive. All donors are displayed against a common lower limit of quantification (29 pg/ml); **n.d.**, not detected; hatched, not measured. One value exceeded the upper limit and is shown as a lower bound.

Because CD63 also reports activation in other myeloid cells, we examined monocytes and neutrophils in the same samples. Binary ICs activated monocytes moderately but neutrophils only marginally, whereas ternary ICs activated both populations (Fig. 2e,f). Monocytes responded across the full concentration range, whereas neutrophil activation was confined to the highest concentration tested. We next stimulated isolated CD14 monocytes overnight with binary ICs, which contain no recombinant allergen and therefore no residual endotoxin. Two populations were distinguished by side scatter, and phenotypic changes were largely confined to the SSC^hi^ population (Fig. 2g, Fig. S4). We monitored CD86, CD11c, HLA-DR, CD83 and CD1a; baseline CD1a expression was low in all donors (Fig. S4). Responses varied between donors, but ligelizumab ICs tended to increase the proportion of marker-positive cells relative to omalizumab ICs (Fig. 2h). Ligelizumab ICs significantly up-regulated CD86 MFI compared with omalizumab ICs (p = 0.011) and gave higher CD11c and HLA-DR in all four donors, although these did not reach significance (Fig. 2i); CD83 MFI did not differ between the two antibodies (Fig. S4). Cytokine output differed between donors in both magnitude and direction (Fig. 2j). In two donors, both antibodies induced TNF-α, IL-1β and IL-6, with omalizumab giving the higher TNF-α and IL-1β values; in the other two, omalizumab induced no detectable TNF-α or IL-1β whereas ligelizumab induced a strong response. MCP-1 was higher with ligelizumab ICs in all donors measured.

Together, these cell-type- and antibody-dependent activation patterns indicated that engagement of any single Fc receptor cannot fully explain IC activity. We therefore asked whether omalizumab- and ligelizumab-containing ICs differ in the balance of Fcε- and Fcγ-receptor interactions they support, and whether that balance itself changes with allergen incorporation.

### Allergen incorporation reverses Fc receptor engagement by omalizumab and ligelizumab

Directly resolving this receptor-balance hypothesis in primary cells is challenging because multiple Fc receptors are co-expressed, endogenous IgE and IgG can occupy them, receptor expression varies between donors and blocking reagents perturb several pathways. Primary-cell assays therefore report the integrated biological consequence of IC recognition but cannot readily identify the receptor-interaction landscape that produced it.

We therefore used the FcγR/FcεR panel of focal molography, a multiplexed, real-time, label-free interaction analysis method as a direct molecular test of whether antibody identity and allergen incorporation rewire Fc receptor engagement. The novel seven-receptor panel comprised FcγRI, FcγRIIa, FcγRIIb, FcγRIIIa, FcγRIIIb, FcεRI and FcεRII (CD23), together with an in-chip negative control sensor. Because the same preformed IC injection was measured simultaneously across all receptor sensors, the format enabled direct real-time comparison of free IgE, therapeutic antibodies and defined binary and ternary ICs (Fig. 3a–d). Representative sensorgrams and per-analyte fit summaries are shown in Supplementary Fig. S6.

**Figure 3.**
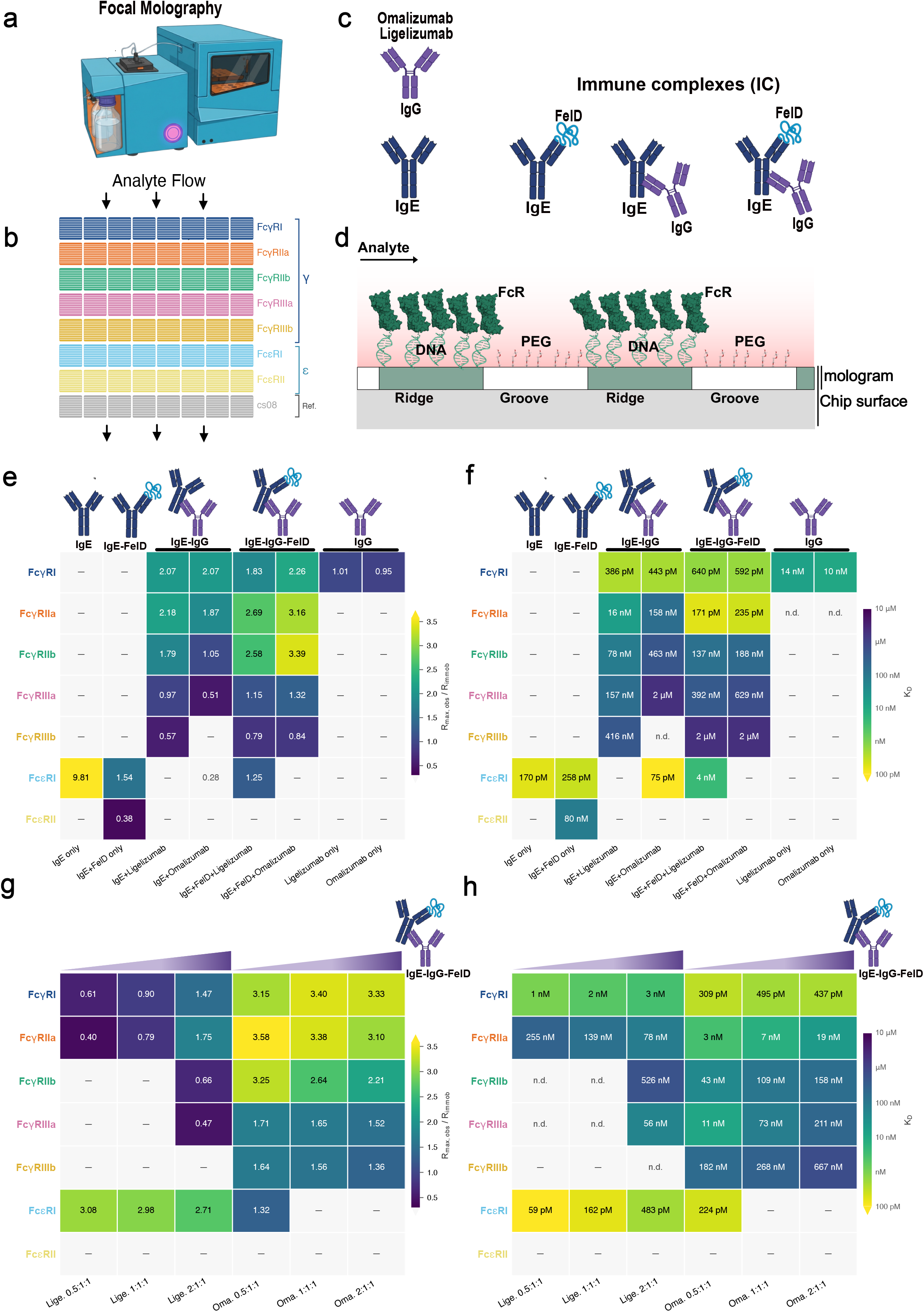
| Allergen incorporation reverses Fc receptor engagement by omalizumab and ligelizumab. **a,** The Lino Biotech MACS Matchmaker (Lino Biotech instrument). **b,** Layout of the multiplex mologram chip showing replicate sensors functionalized with FcγRI, FcγRIIa, FcγRIIb, FcγRIIIa, FcγRIIIb, FcεRI, FcεRII/CD23, and an in-chip negative control sensor column (cs08). Arrows indicate analyte flow across the receptor panel. **c,** Schematic of the analytes and IC species profiled. **d,** Schematic of the mologram chip surface **e,** Normalized maximal binding response (R_max_/R_immo_) across the receptor panel. **f,** Apparent equilibrium dissociation constants (K_D_). Lower K_D_ values indicate stronger binding. **g,** normalized maximal binding response (R_max_/R_immo_) for ternary ICs assembled at anti-IgE:IgE:Fel d 1 molar ratios of 0.5:1:1, 1:1:1, and 2:1:1, with IgE and Fel d 1 held constant. **h,** Apparent K_D_ values for the same stoichiometric series. Cells marked with “–” indicate no measurable binding above the analysis threshold. n.d., not determinable from kinetic fitting. Signal is reported as Coherent Mass Density (pg/mm²). Data shown are for IgE clone F127; equivalent binary and ternary IC profiling for IgE clones A044 and G078 is provided in Supplementary Figs. S9 and S10.

Monomeric IgE bound strongly to FcεRI and showed no detectable interaction with Fcγ receptors, whereas free omalizumab and ligelizumab engaged FcγRI without detectable binding to Fcε or low-affinity Fcγ receptors (Fig. 3e, f). Formation of IgE–Fel d 1 complexes increased binding to FcεRII (CD23). Binary IC formation then created a new receptor state. Both IgE–IgG complexes broadened engagement across low-affinity FcγRs while losing Fcε-receptor binding, but ligelizumab-containing binary ICs engaged FcγRIIa/b and FcγRIIIa/b more strongly than omalizumab-containing ICs (K_D_; Fig. 3f), with comparable FcγRI binding. Ligelizumab also suppressed FcεRI engagement more completely: omalizumab-containing ICs retained a small, high-affinity FcεRI-binding population (R_max_/R_immo_ 0.28; K_D_ 75 pM), whereas ligelizumab-containing ICs showed no detectable FcεRI binding (Fig. 3e, f).

Allergen incorporation was the central surprise: it reversed the omalizumab–ligelizumab receptor hierarchy. In ternary ICs, ligelizumab lost its advantage in suppressing FcεRI, whereas omalizumab more effectively prevented FcεRI engagement. Simultaneously, omalizumab gained FcγR binding ability relative to ligelizumab. At 1:1:1 stoichiometry, this inversion was also observed with additional Fel d 1-specific IgE clones (Supplementary Figs. S9 and S10).

When anti-IgE was varied 0.5-, 1-, or 2-fold relative to fixed IgE and Fel d 1, omalizumab-containing ternary ICs maintained strong normalized responses across FcγRI, FcγRIIa/b and FcγRIIIa/b, whereas FcεRI binding was detectable only at 0.5-fold omalizumab and fell below the measurement threshold at equimolar and twofold antibody (Fig. 3g; Supplementary Fig. S7). Ligelizumab-containing ternary ICs showed the opposite balance: FcγR responses remained lower and only became measurable for several receptors at the highest antibody ratio, while FcεRI engagement persisted across all ratios. Apparent dissociation constants mirrored this separation: omalizumab complexes showed subnanomolar-to-nanomolar FcγRI/IIa interactions, whereas ligelizumab complexes retained high-affinity FcεRI binding throughout the series (Fig. 3h).

These data establish a broader principle: anti-IgE function is state-dependent and cannot be predicted from free-IgE affinity alone. The same therapeutic antibody adopts a different Fc-receptor interaction profile when IgE shifts from a binary anti-IgE complex to an allergen-containing ternary complex. We therefore next asked whether this qualitative receptor inversion produces a meaningful quantitative consequence for FcεRI neutralization.

### Ligelizumab requires a more than 20-fold higher concentration than Omalizumab to neutralize FcεRI engagement by allergen-bound IgE

We next asked whether the qualitative receptor inversion translated into a meaningful quantitative difference in neutralizing potency. To test this, we assembled ternary IgE– Fel d 1 ICs with increasing concentrations of omalizumab or ligelizumab and determined the concentration required to inhibit 50% of FcεRI engagement (IC□□) by multiplex focal molography (Fig. 4a–d and Supplementary Fig. S8).

**Figure 4.**
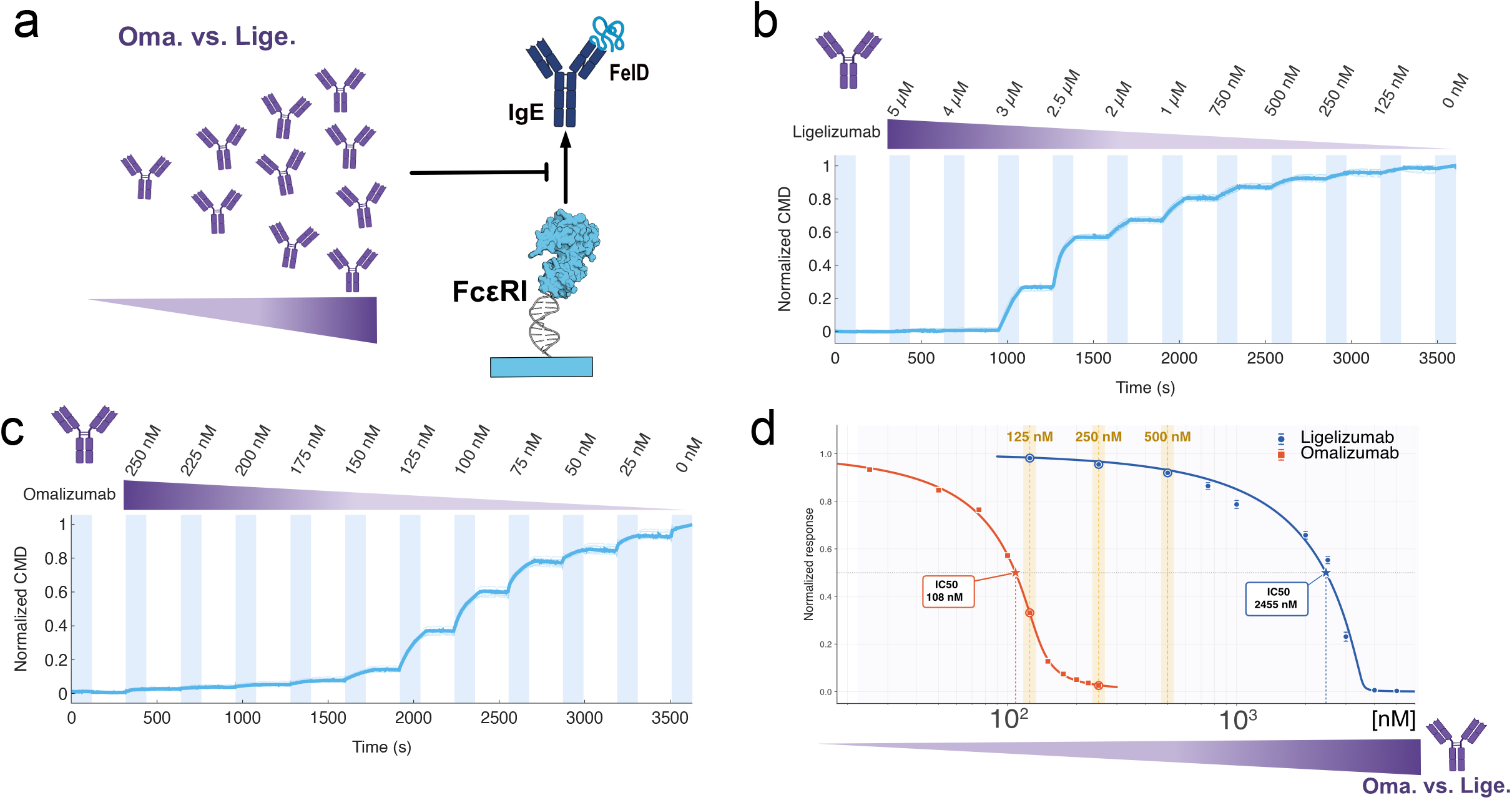
| Ligelizumab requires a more than 20-fold higher concentration than Omalizumab to neutralize FcεRI engagement by allergen-bound IgE. Ternary ICs were assembled with a titration of omalizumab or ligelizumab, and FcεRI engagement was quantified by multiplex focal molography. **a**, Schematic of the assay: increasing concentrations of omalizumab or ligelizumab were premixed with IgE and Fel d 1, and residual FcεRI engagement of the resulting ternary complex was measured on the mologram surface. **b–c**, Single-cycle sensorgrams (coherent mass density versus time) recorded on the FcεRI column across the ligelizumab (**b**) and omalizumab (**c**) concentration series; faint traces show the eight individual replicate mologram spots and the bold trace their median, and shaded bands mark the successive association steps of the concentration titration. **d**, Normalized FcεRI engagement as a function of anti-IgE concentration. Points show the median across eight replicate mologram spots within one chip, and error bars show the corresponding SD. Curves were fitted using the mechanistic titration model described in Methods. The reported bootstrap intervals quantify within-chip technical precision and do not represent between-experiment reproducibility. Sensorgrams of the individual titration series are provided in Supplementary Fig. S8; the omalizumab and ligelizumab titrations were performed on separate chips.

Omalizumab achieved half-maximal FcεRI inhibition at approximately 108 nM (95% CI, 107–109 nM), whereas ligelizumab required approximately 2,455 nM (95% CI, 2,382– 2,530 nM), an approximately 23-fold higher concentration. For comparison, a simple stoichiometric model predicts that binding 50% of the 250 nM IgE would require 62.5– 125 nM anti-IgE, depending on whether each IgG engages two IgE molecules or one. Omalizumab’s IC□□ fell within this range, whereas ligelizumab’s was approximately 20–39-fold higher. Although this comparison assumes that IgE binding translates directly into inhibition of receptor engagement, it provides a useful benchmark for the observed difference in neutralizing potency.

Thus, the allergen-driven qualitative receptor inversion culminated in an approximately 23-fold separation in FcεRI-neutralizing potency: omalizumab approached stoichiometric neutralization, whereas ligelizumab did not.

## Discussion

We show that the functional properties of therapeutic anti-IgE antibodies depend on the composition of the ICs they form. In primary leukocytes, we observe antibody- and composition-dependent patterns of cellular association and activation. At the molecular level, ligelizumab more effectively neutralized FcεRI interactions in binary IgE–IgG complexes and generated a stronger low-affinity Fcγ receptor-binding signature than omalizumab. Incorporation of Fel d 1 into ternary ICs reversed this relationship: omalizumab more effectively neutralized FcεRI engagement and produced stronger low-affinity Fcγ binding. Thus, the functional ranking of omalizumab and ligelizumab inferred from free-IgE measurements does not predict their behaviour in allergen-containing ICs.

Therapeutic anti-IgE antibodies are primarily evaluated by their ability to neutralize IgE– FcεRI interactions. However, IgE–IgG ICs are well documented in omalizumab-treated patients and contribute to increases in total serum IgE following therapy^22^. Our findings support a broader view in which the functional properties of ICs emerge from their composition rather than from any single antigen–antibody or receptor interaction. Therapeutic anti-IgE antibodies provide a particularly clear example of this principle: changing the anti-IgE antibody or incorporating allergen reshaped the balance of Fcε and Fcγ receptor interactions and produced distinct functional consequences.

In our panel, CD23 engagement required complexed rather than free IgE: monomeric IgE showed no detectable CD23 binding, whereas IgE–Fel d 1 complexes bound CD23 (Fig. 3e, f). Previous work has reported that omalizumab inhibits IgE-IC binding to CD23 more effectively than ligelizumab, which we here also confirm (Fig. S8)^10,11,13^. Our observations are likewise consistent with the distinct IgE epitopes for omalizumab and ligelizumab^11^. One possibility is that allergen binding alters the conformation, accessibility, or orientation of the IgE Fc region, thereby changing receptor engagement. Functionally, our findings point to additional IC-dependent mechanisms that distinguish the two antibodies. The targeting of allergen towards Fcγ receptor-expressing leukocytes may provide an additional route through which therapeutic anti-IgE antibodies influence immune responses. Monocytes may be a cell type of particular relevance, as they are implicated in allergic inflammation and asthma, yet remain relatively understudied in the context of therapeutic anti-IgE responses^23^.

Thus, our findings provide several possible mechanistic explanations for the apparent ligelizumab paradox, whereby its enhanced ability to neutralize IgE–FcεRI interactions did not translate into clinical superiority over omalizumab in allergic disease:

First, allergen-containing ligelizumab ICs may retain a higher capacity to activate pre-sensitized basophils, as observed in our experiments, in which responses were most evident in donors with pre-existing Fel d 1-specific IgE. A similar mechanism could operate in allergen-sensitized mast cells. Thus, the functional consequences of anti-IgE therapy may depend both on the pre-existing sensitization state of effector cells and on the properties of ICs formed during ongoing allergen exposure.

Second, altered engagement and activation of FcγR-expressing effector cells, such as monocytes or neutrophils, could contribute to differences in the biological activity of ligelizumab-containing ICs. In our setting, binary ICs activated monocytes but only marginally activated neutrophils, whereas ternary ICs activated both. A potential explanation is that binary ICs engaged FcγRI and FcγRII with higher apparent affinity than FcγRIII. In monocytes, binary ICs further modulated maturation markers and cytokine responses in an antibody-dependent manner.

While free-IgE neutralization remains a central mechanism underlying the clinical efficacy of anti-IgE therapy, our findings suggest that profiling IC signatures may provide a more comprehensive understanding of their biological activity. More broadly, these observations raise the possibility that therapeutic anti-IgE antibodies must strike a balance between efficient neutralization of free IgE or IgE-allergen complexes, and the binding and/or activation of Fcγ receptors. The relative importance of these pathways may also vary with the respective disease context.

Methodologically, this work establishes multiplex focal molography for the real-time profiling of IC interactions across the human Fc receptor landscape. Unlike conventional biophysical assays or reporter-cell systems that interrogate individual receptors, multiplex profiling captures the broader receptor interaction profile of higher-order ICs and provides a molecular view of their potential cellular interactions. Because IC formation is central to many antibody therapeutics, this approach may prove broadly useful for mechanistic studies and therapeutic antibody development.

Our study has limitations. ICs are concentration-dependent equilibrium assemblies rather than defined molecular species, and no standardised preparations exist against which they can be benchmarked; ours are defined only by input stoichiometry. Because these complexes are heterogeneous and multivalent, no single kinetic descriptor captures their behaviour: maximal binding response and apparent affinity reflect complex size and receptor valency differently and need not rank analytes identically. Biologically, the recombinant allergen and monoclonal IgE system is deliberately reductionist, and basophil data come from few, highly variable donors. However, the FcεRI reversal held in all donors and across three IgE clones (Supplementary Figs. S9 and S10). Whether this extends to other allergen specificities, polyclonal patient IgE, larger patient cohorts and natural exposure remains open.

Together, our findings demonstrate that therapeutic anti-IgE antibodies cannot be fully understood through their interaction with free IgE alone. Instead, the ICs generated by these antibodies reshape Fc receptor engagement, cellular association and activation. We therefore propose that future anti-IgE therapeutics should be evaluated not only for free IgE and FcεRI neutralization, but also for the functional IC signatures they generate.

## Methods

### Experimental design

To determine how IC composition shapes Fc receptor engagement and cellular function, we generated binary IgE–IgG and ternary IgE–IgG–Fel d 1 complexes containing omalizumab or ligelizumab at defined stoichiometry. We assessed their association and activation in primary human leukocytes, examined cytokine responses in purified monocytes and profiled their interactions with major Fcε and Fcγ receptors using multiplex focal molography.

### Single-cell transcriptomic analysis of Fc receptor expression

Publicly available single-cell RNA sequencing data from human whole blood (10x Genomics; *Aggregate of Human Whole Leukocytes Isolated from Fixed Whole Blood of a Diseased Donor and a Healthy Donor*, Single Cell Fixed RNA Profiling Dataset analysed by Cell Ranger v8.0.0, March 27, 2024) were analysed using Seurat (v5) in R. The filtered feature-barcode matrix was log-normalized, variable genes were identified, and cells were clustered using the Louvain algorithm using a low resolution of 0.03 to broadly cluster cell types, following principal component analysis and nearest-neighbour graph construction. Cell populations were annotated manually based on established lineage-specific marker genes and grouped into the major immune cell populations analysed by flow cytometry. Fc receptor gene expression was visualized using UMAP feature plots and dot plots.

### Recombinant proteins and IC generation

Recombinant dimeric Fel d 1 and three independent human Fel d 1-specific IgE clones (G078, F127, A044) were generated and characterized as previously described20,24. Unless otherwise indicated, experiments were performed using IgE clone F127. Binary and ternary IC profiling by focal molography was additionally performed with IgE clones A044 and G078 (Supplementary Figs. S9 and S10). For cellular binding assays, Fel d 1 was fluorescently labelled using Alexa Fluor 488 or 647 labelling kits (Thermo Fisher Scientific). Omalizumab and ligelizumab were kindly provided by Dr. Christoph Heusser (Formerly Novartis Pharma, Basel, Switzerland). ICs were assembled by mixing components in the order IgE → Fel d 1 → anti-IgE and incubating for 1 h at room temperature before use. Unless otherwise indicated, binary complexes were assembled at 1:1 stoichiometry; and ternary complexes were assembled at 1:1:1 stoichiometry.

### Human donors

The study was approved by the local ethics committee (Kantonale Ethikkommission Bern, references 2020-00699, 2026-00605). Peripheral venous blood was collected from healthy adult volunteer donors. Specific IgE against recombinant Fel d 1 (e94/rFel d 1) was measured by ImmunoCAP (Thermo Fisher Scientific). Whole blood was subjected to erythrocyte lysis using ACK buffer. Alternatively, PBMCs were isolated by Ficoll-Paque density-gradient centrifugation. Leukocytes were resuspended in complete RPMI 1640 medium with 10% FBS and plated at 1–2 × 10□ cells per well. For experiments using purified monocytes, CD14 cells were isolated by positive immunomagnetic selection (CD14 MicroBeads, Miltenyi Biotec, cat. 130-050-201; MS Column, cat. 130-042-201) following erythrocyte lysis and seeded at 9.5 × 10□ cells per well.

### Statistical analysis of human data

Statistical analyses of cellular data were performed in GraphPad Prism (version 11; GraphPad Software). *Data presentation.* Where no symbols are shown, data are mean ± SEM of technical replicates from a single donor, representative of two independent experiments performed with that donor. Where symbols are shown, each represents one donor; technical replicates were averaged within each donor before analysis. Donor numbers are given in the figure legends. *Area under the curve.* For basophil activation, AUC was calculated per donor and condition from the concentration–response curve of % CD63□ cells against Fel d 1 concentration (0–200 nM, linear scale, trapezoidal rule). Because the concentration series is logarithmically spaced, the 20–200 nM interval contributes most of the integrated area. Analyses used untransformed AUC. *Hypothesis testing.* Allergen association across four concentrations (Fig. 1h) was analysed by two-way repeated-measures ANOVA matched on both factors, with antibody (omalizumab, ligelizumab) and concentration as within-donor factors (n = 3 donors), Geisser– Greenhouse correction, and Šídák’s multiple comparisons test comparing the two antibodies at each concentration. Basophil AUC (Fig. 2d) was analysed by two-way repeated-measures ANOVA with antibody (omalizumab, ligelizumab) and IC composition (binary, ternary) as within-donor factors (n = 5 donors). The reported significance refers to the antibody × IC interaction. Simple effects within each IC class were not significant (binary p = 0.16, ternary p = 0.060), and neither main effect reached significance, as expected where the effect reverses direction between IC classes. Monocyte and neutrophil CD63 responses (Fig. 2e, f) were compared against the IgE–Fel d 1 reference condition by two-way repeated-measures ANOVA followed by Dunnett’s multiple comparisons test (n = 4 donors). Monocyte maturation markers (Fig. 2i) were compared between omalizumab- and ligelizumab-containing ICs by two-tailed paired t-tests (n = 4 donors. *Descriptive data.* The following were not statistically tested: single-donor binding experiments (Fig. 1e, f); allergen association at 37 °C (Fig. 1g); basophil concentration–response curves and per-donor AUC values (Fig. 2b, c); and cytokine data (Fig. 2j, Fig. S5). *Significance.* A two-sided p value < 0.05 was considered statistically significant. *p < 0.05, **p < 0.01, ***p < 0.001, ****p < 0.0001.

### In vitro assays with primary immune cells

For binding assays, the cells were incubated with complexes consisting of equimolar IgE, recombinant Fel d 1, omalizumab, ligelizumab (0.2, 2, 20, or 200 nM) for 30 min at 4°C or 37°C, stained with fluorochrome-conjugated antibodies (Table 1), and analysed on a CytoFLEX S flow cytometer (Beckman Coulter). Dead cells were excluded using Fixable Violet Viability (Thermo Fisher Scientific). Binding was quantified as the frequency of positive cells or geometric mean fluorescence intensity (gMFI). For overnight stimulation assays, FcγRII/III receptors were blocked prior to stimulation by addition of human FcR Blocking Reagent (BD Biosciences, cat. 564220; 1:100 final dilution) for 15 min at room temperature without subsequent washing. The plates were incubated o/n at 37°C, 5% CO□. After incubation, supernatants were collected and stored at −20°C; cytokine concentrations were measured using the LEGENDplex™ Human Essential Immune Response Panel (13-plex, BioLegend, cat. 740929).

**Table 1:**

| Antibody | Conjugate | Clone |  | Cat | RRID |
|---|---|---|---|---|---|
| Fcer1a | FITC | AER-37 (CRA1) | Thermo Fisher Scientific | 11-5899-41 | AB_10732343 |
| CD23 | PE | EBVCS-5 | BioLegend | 338508 | AB_1279177 |
| CD16 | PE | B73.1 | BD Biosciences | 561313 | AB_10643606 |
| CD32 | PE | FLI8.26 | BD Biosciences | 550586 | AB_393766 |
| CD64 | PE | 10.1 | BioLegend | 305008 | AB_314492 |
| CD63 | FITC | H5C6 | BD Biosciences | 557288 | AB_396617 |
| CD3 | PerCP-Cy5.5 | UCHT1 | BioLegend | 300430 | AB_893299 |
| CD123 | PE | 6H6 | BD Biosciences | 555644 | AB_396001 |
| CD19 | PE-Cy7 | HIB19 | Thermo Fisher Scientific | 25-0199-42 | AB_1582278 |
| CD11b | APC-Cy7 | M1/70 | BioLegend | 101226 | AB_830642 |
| CD11c | BV480 | 3.9 | Thermo Fisher Scientific | 414-0116-42 | AB_3714955 |
| CD86 | FITC | 2331 (FUN-1) | BD Biosciences | 555657 | AB_396012 |
| HLA-DR | PerCP | L243 | BioLegend | 307628 | AB_893566 |
| CD1a | Cardinal Red | HI149 | Thermo Fisher Scientific | CR-65172 | AB_3084971 |
| CD83 | APC-eFluor 780 | HB15e | Thermo Fisher Scientific | 47-0839-42 | AB_2848359 |

### Flow cytometry gating strategy

Flow cytometry data were analysed using a sequential gating strategy. Fluorescence-minus-one controls were used where appropriate to define positive populations. Debris and low-quality events were excluded based on FSC/SSC characteristics, followed by selection of single cells using FSC-A versus FSC-H. Viable leukocytes were identified by exclusion of dead cells using the viability dye. Major leukocyte populations were subsequently identified using lineage-associated surface markers, including T cells (CD3□), B cells (CD19□), basophils (CD123□CD11b□), pDCs (CD123□CD11b□), monocytes (CD14□CD11b□), and neutrophils based on FSC/SSC properties and CD11b expression. CD63 expression was quantified within the respective basophil (CD123□CD11b□), monocyte, or neutrophil gates. For purified monocyte experiments, two populations were distinguished based on side-scatter characteristics (SSC^hi^ and SSC^lo^), and expression of CD83, CD86, HLA-DR, CD11c, and CD1a was quantified within the respective populations. Identical gating criteria were applied across experimental conditions within each donor.

### Focal molography-based Fc receptor profiling

Fc receptor interactions were analysed using a multiplex focal molography platform on the MACS Matchmaker. The signal recorded is the Coherent Mass Density at the focal spot of each mologram; this is a real-time, label-free measure of analyte mass bound to the immobilized ligand and is distinct from refractive-index-based SPR/BLI readouts. 8-PLEX-PEG sensor chips were used. Both running and sample buffers were PBS pH 7.4 supplemented with 0.05% (v/v) Tween-20 (PBS-T). Chip regeneration between runs was performed with 3 M guanidine hydrochloride / 125 mM NaOH (0.5 min at 400 µL/min) bracketed by PBS-T washes (3 min at 300 µL/min).

### Receptor immobilization

Recombinant human Fc receptors FcγRI, FcγRIIa, FcγRIIb, FcγRIIIa, FcγRIIIb, FcεRIα, and FcεRII/CD23 (Gamma Proteins, AcroBiosystems) were pre-conjugated to spot-specific oligonucleotides (cs01-cs07) and captured on the molographic sensor surface by DNA-directed immobilization at 200 nM each in 150 µL PBS-T with a reference spot (cs08) (Biomers). Receptor mixtures were injected at 10 µL/min for 10 min (single-analyte assays) or 5 min (binary and ternary assays).

### Single-cycle kinetics (SCK)

Single-cycle kinetics (SCK) measurements were performed following a PBST/regeneration/PBST baseline. Unless otherwise stated, samples were injected as concentration series with 2 min association at 30 µL/min and 2 min dissociation at 200 µL/min between consecutive injections, followed by a 30 min final dissociation at 200 µL/min. Surfaces were regenerated between cycles using 3 M guanidine HCl/125 mM NaOH. For individual analytes, IgE, omalizumab, or ligelizumab were injected over a concentration range of 0–1 µM. For binary IC, IgE + Fel d 1 or IgE + anti-IgE were premixed at a 1:1 stoichiometry and incubated for 1 h at 25 °C with shaking at 500 rpm before injection. SCK concentration ranges were 0–100 nM for IgE + Fel d 1 and 0–500 nM for IgE + anti-IgE. For ternary IC, IgE, Fel d 1, and anti-IgE were premixed at a 1:1:1 stoichiometry in the order IgE → Fel d 1 → anti-IgE and incubated for 1 h at 25 °C with shaking at 500 rpm before injection. Complexes were analysed over an SCK concentration range of 0–750 nM. For stoichiometry titrations, IgE (clone F127) and Fel d 1 were each held at 250 nM, while anti-IgE (omalizumab or ligelizumab) was varied between 125, 250, and 500 nM. Complexes were prepared using the same pipetting order and pre-incubation conditions described above. For these measurements, the final dissociation phase was extended to 60 min.

### Kinetic data evaluation and preprocessing

Single-cycle kinetics molograms were evaluated with a custom Python pipeline wrapped in an interactive Streamlit interface. Each chip carried seven receptor columns (C1–C7), each represented by 8 replicate mologram spots, together with one in-chip reference column (C8). The fitting unit was the individual sensorgram (one per spot per cycle), and column-level summary statistics were obtained by aggregation across replicate sensorgrams within each receptor column. Raw time traces (Coherent Mass Density, pg/mm²) were imported per spot and grouped by column. The SCK association block was localized automatically from injection metadata, and analyte identity was extracted from the sample record. Reference subtraction was performed point-by-point using the C8 reference column, followed by subtraction of any blank-injection group corresponding to the 0 M cycle when present. For each association injection, endpoint response was defined as the mean signal over the last 20% of the association phase minus a baseline calculated as the mean of the first 5% of the first association phase. Columns were retained for fitting only if their maximum endpoint response reached at least 20% of the highest-responding column in the run; columns below this cross-column threshold were classified as no binding and excluded. Within retained columns, the lowest concentration steps were trimmed adaptively until endpoint signal reached at least 20% of that column’s own maximum response, and a minimum of three concentration steps was required for fitting.

### Off-rate estimation

For each retained column, the highest-concentration dissociation phase was first evaluated per sensorgram with a single-exponential decay model to assess dissociation behaviour. Columns showing evidence of a non-dissociable residual were re-estimated using an exponential decay with baseline term. Off-rate estimates were then refined on adaptive short dissociation windows to reduce baseline bias. When the predicted dissociation over the available window was below 5%, the off-rate was replaced by the corresponding minimum detectable value, and a hard lower floor of k_off_ ≥ 2.7 × 10□□s^□^¹ was enforced. Sensorgrams affected by these detectability limits were flagged in the output. For the optional long-dissociation analysis path, two per-column models were available: single-exponential decay and single-exponential decay with a non-dissociable baseline. Degenerate fits were identified using pre-specified criteria that combined implausibly fast off-rates with near-perfect nominal decay or extremely short apparent half-lives relative to the fit window. When such cases occurred, the dissociation tail was re-fitted after exclusion of the earliest part of the trace; unresolved cases were classified as below detectability, capped at the minimum detectable off-rate, and excluded from the column median.

### Global kinetic fitting

For each retained ligand, all selected concentration steps were fitted jointly per sensorgram using either a 1:1 Langmuir model or a 1:1 Langmuir model with a shared non-dissociable residual fraction. Baseline offsets were removed directly on trimmed sensorgrams using the last 20% of the first dissociation phase, or when appropriate the first 5% of the first association phase, as reference. Fitting was performed in log-space on the rate constants, and sensorgrams were internally scaled to unit amplitude during optimization before fitted R_max_ values were transformed back to physical units. Automatic adjustment of the initial response term was enabled to absorb residual baseline drift between injections. Two off-rate source modes were used: a global-fit mode in which both k_on_ and k_off_ were optimized directly from the SCK traces, and a long-dissociation mode in which k_off_ from the long-dissociation pre-fit was fixed per sensorgram while k_on_ and R_max_ were optimized. For fits with free k_off_, an internal consistency check compared the fitted value with the long-dissociation estimate and replaced the fitted value when the predicted decay across the dissociation window fell below the detectability threshold.

### Result aggregation and quality control

For every sensorgram, the fit returned k_on_, k_off_, K_D_ = k_off_/k_on_, R_max_, and reduced χ². Column-level reduced χ² values were aggregated as the median across replicate sensorgrams. Individual sensorgram fits were flagged as poor when reduced χ² exceeded 10^□^², and a receptor column was classified as not determinable when more than 50% of its sensorgrams were flagged. K_D_ values derived from off-rates capped at the detectability floor were marked to indicate that k_off_ was not fully constrained by the data. Inter-replicate outliers within a column were identified in log-space using a median absolute deviation rule. Final exports included per-sensorgram fit results and column-level summary statistics, including K_D_ geometric means, median k_on_ and k_off_ values, and inter-replicate spread.

### Determination of inhibitory potency (IC□□)

#### Assay and signal acquisition

Inhibition of Fel d 1-specific IgE (Fel d 1–IgE-F127) binding to immobilized FcεRI was measured by focal molography on a MACS® Matchmaker instrument. The reported signal is the normalized coherent mass density (CMD), acquired simultaneously from eight replicate mologram spots (n = 8) per chip. A descending single-cycle titration series of anti-IgE antibodies (omalizumab or ligelizumab) was pre-incubated with a fixed concentration of Fel d 1–IgE-F127 (IgE at 250 nM) and injected over the sensor surface (sensorgrams in Supplementary Fig. S8). Buffer-reference cycles were interleaved throughout the titration and excluded from the analysis. *Response extraction.* For each spot and each concentration step, FcεRI surface occupancy was derived from the normalized CMD and scaled relative to the uninhibited control, such that the 0 nM anti-IgE condition corresponds to an occupancy of 1.0. Occupancy was summarized across the eight replicate spots as the median with the corresponding standard deviation at each concentration. *Dose–response fitting.* The anti-IgE antibodies bind IgE with an affinity substantially higher than the IgE concentration employed, so the titration operates in the tight-binding, stoichiometric regime. This regime generates a characteristically asymmetric dose–response — a shallow plateau at low inhibitor concentrations followed by an abrupt transition — that a symmetric four-parameter logistic (Hill) function cannot reproduce without invoking a physically unjustified Hill coefficient. Occupancy data were therefore fitted with a mechanistic titration model that couples a quadratic (Morrison) binding equilibrium for the antibody– IgE interaction to a saturable Langmuir readout of the residual free IgE at the sensor surface:

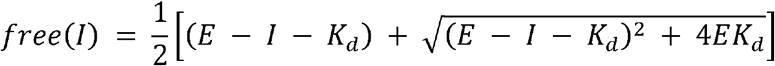

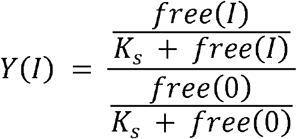

where I denotes the anti-IgE concentration, E the binding-competent IgE concentration, K_D_ the apparent antibody–IgE dissociation constant, and Ks the surface dissociation constant for IgE. Parameters were estimated by non-linear least squares (curve_fit, SciPy) with weighting by the inter-spot standard deviation at each concentration. IC□□ was determined numerically as the concentration satisfying Y = 0.5, using Brent’s root-finding method.

#### Uncertainty estimation

Confidence intervals were derived by a cluster bootstrap in which the eight replicate spots were resampled with replacement over 3,000 iterations. For each resample the per-concentration median and standard deviation were recomputed and the mechanistic model refitted; the 2.5th and 97.5th percentiles of the resulting IC□□ distribution define the reported 95% confidence interval.

## Data and code availability

Focal Molography and Flow Cytometry source data and analysis code are deposited at Zenodo DOI 10.5281/zenodo.23973836. Further information is available from the corresponding author on request.

## Author contributions

P.E. conceptualized the study. P.E., and M.F.B. acquired research funding and ethical permission. P.E., and M.F.B. supervised the study. A.F supervised the focal molography experiments. M.M.N. established, designed, and performed the flow cytometry assays in primary immune cells, and analysed the corresponding data. S.N. and A.G.G. performed focal molography experiments and analysed the data. P.E., M.M.N., and A.G.G. visualized the data and wrote the manuscript. All authors discussed the results, critically reviewed and edited the manuscript, and approved the final version.

## Acknowledgements

We thank Dr. Christoph Heusser (formerly Novartis Pharma AG, Basel, Switzerland) and Prof. Dr. Monique Vogel (University of Bern, Bern, Switzerland) for initiating this project. We thank Marc Huldi (formerly University of Bern, Bern, Switzerland) for contributions to the early experimental work that helped shape its subsequent direction. We thank Dr. Ulf Gehrmann (Novartis Pharma AG, Basel, Switzerland) for instructive discussions on the resulting data. We thank Marianne Zwicker and Sandra Nansoz for technical assistance. We thank Dr. Andrea Karolin (Zentrum für Labormedizin, Inselspital Bern) for specific IgE measurements. We thank the donors who consented to provide blood for this study and the entire staff of the Department of Rheumatology and Immunology, Inselspital Bern, for technical assistance with donor recruitment and sample collection.

